# Rhamnose Polysaccharide-Decorated Outer Membrane Vesicles as a Vaccine Candidate Targeting Group A Streptococcus from *Streptococcus pyogenes* and *Streptococcus dysgalactiae* subsp. *equisimilis*

**DOI:** 10.1101/2024.08.23.609347

**Authors:** Sowmya Ajay Castro, Sarah Thomson, Helen Alexandra Shaw, Azul Zorzoli, Benjamin H Meyer, Mark Reglinski, Helge C. Dorfmueller

## Abstract

Group A Streptococcus (Strep A) cause a wide range of human-exclusive infections, annually killing more than 500,000 people. Antibiotic resistance incidence of invasive Strep A tripled in the past decade and emphasises the need to develop a universal Strep A vaccine. In this study, we developed recombinant rhamnose polysaccharides (RhaPS), a validated universal Strep A vaccine candidate, presented on *E. coli* outer membrane vesicles (OMVs). We investigated OMV-RhaPS for their immunogenicity in the mouse and rabbit models. Through flow cytometry, ELISA, Luminex assays, and immunofluorescence microscopy, we demonstrated that RhaPS-specific antibodies recognise Strep A strains via the Group A Carbohydrate (GAC) in *S. pyogenes* and the newly emerged *S. dysgalactiae* subsp. *equisimilis*. Elevated IL-17a levels from RhaPS-OMV-immunised splenocytes indicate the RhaPS-specific stimulation of long-term memory immune cells. We are the first to report the efficacy and potency of recombinant produced RhaPS inducing humoral-mediated immune responses and triggering antibodies that recognise Strep A bacteria.

## Introduction

*Streptococcus pyogenes*, frequently described as Group A Streptococcus (Strep A), is a beta-haemolytic Gram-positive human exclusive bacterium causing a wide range of illnesses in both healthy and immunocompromised individuals. Breaching of immune barriers by Strep A leads to acute suppurative effects in the skin and pharyngeal epithelia, causing localised skin infections, tonsillitis and enlarged lymph nodes in the throat [1]. Untreated Strep A can lead to life-threatening complications such as streptococcal toxic shock syndrome, necrotising fasciitis and, importantly, post-streptococcal autoimmune sequelae [2]. One of these major autoimmune sequelae is rheumatic heart disease, which kills >300,000 patients worldwide [1, 3]. Incidence of antibiotic resistance has tripled in the past decade, and the common penicillin-based antibiotics only eradicate ∼65% of all tonsilitis cases caused by Strep A infections [4, 5].

Identification of Strep A is based on the detection of its main surface antigen, the Group A Carbohydrate [GAC]. This consists of a polyrhamnose (RhaPS) backbone (→3)α-Rha(1→2)α-Rha(1→) with an immunodominant N-acetylglucosamine (GlcNAc) side chain present on every α-1,2-linked rhamnose [6-8]. Recent studies have reported the presence of negatively charged glycerol-phosphate onto approximately every fourth GlcNAc sidechain [9]. Earlier research proposed GAC as a potential antigen due to i) its abundant presence in the cell wall contributing ∼50% of Strep A by weight, ii) its conservation across all >200 Strep A serotypes and sequenced Strep A genomes [10], and iii) the absence of cross-reactivity between antibodies raised against endogenously-produced RhaPS and human cardiac antigen [11, 12].

High-titre GAC antibodies have frequently been detected in serum samples, directly correlating with the absence of Strep A in the throats of children [13]. Additionally, anti-GAC antibody titres persist in patients with rheumatic valvular disease [14]. Moreover, a strong body of evidence shows that injecting purified, or synthetic components of GAC conjugated to a carrier protein elicit an immune response that confers protection against multiple Strep A serotypes [13, 15]. Due to its universal conservation and presence in all Strep A isolates, the GAC and the RhaPS backbone are both being explored as vaccine candidates by various research groups. These candidates have shown strong immunogenicity, and recent studies have reported no off-target effects [11, 12, 16] as reviewed in [17].

Outer membrane vesicle (OMV)-based vaccine candidates produced in *E. coli* have been in development for more than ten years and have shown efficacy against several pathogens [18, 19]. Recent efforts have addressed the chemical synthesis of GAC fragments and subsequent engineering of OMVs with the synthesised carbohydrates, revealing that the vesicles provide an additional benefit to the stimulation of carbohydrate specific antibodies when compared with protein conjugates [16].

In this study, we demonstrate that recombinantly produced RhaPS-OMVs can be generated directly from *E. coli* bacteria, thereby eliminating the need for chemical extraction and conjugation steps. We assessed the recombinant glycan structure of RhaPS-OMVs and confirmed, through linkage analysis, that they carry the correct RhaPS structure. This structure is identical to the natively produced RhaPS found in a Strep A mutant strain deficient in the GlcNAc side chain. [11, 12]. We show that *E. coli* RhaPS-OMV are immunogenic in a mouse model; moreover, the stimulated IgG antibodies were able to bind Strep A isolates from *S. pyogenes* and emerging *S. dysgalactiae* subsp. *equisimilis* (SDSE) that carry the *gac* gene cluster and produce the immunogenic GAC surface carbohydrate. Increased IL-17a levels in immunised mice splenocytes suggest the possible stimulation of cellular-mediated immune response in RhaPS-OMV vaccinated animals. We therefore present an alternative route to recombinantly produce a universal Strep A OMV-based vaccine candidate, removing the requirement to produce the Strep A carbohydrate component chemically or extract it chemically or enzymatically from Strep A bacteria and conjugate it to vesicles.

## Results

### Recombinantly produced polyrhamnose-OMVs carry the antigen on Lipid A

The Strep A rhamnose polysaccharide (RhaPS) carbohydrate antigen consists of a linear polymer of rhamnose residues linked by alternating α 1-2 and α 1-3 bonds forming the polysaccharide backbone of Strep A, Group C Streptococci (GCS) and *S. mutans*. RhaPS is also found in newly emerging isolates from *S. dysgalactiae* subsp. *equisimilis* [11, 20-22]. We recombinantly expressed the gene cluster from *S. mutans* (*sccB-G*) and *S. pyogenes* (*gacB-G*) in *E. coli* which encode the biosynthesis machinery for the RhaPS backbone [9, 23]. Our Western blotting analysis of either whole *E. coli* cells or different cellular fractions with Lipid A specific antibodies revealed that the RhaPS produced in these systems are deposited on the outer membrane Lipid A, transferred onto it in the inner membrane *via* the *E. coli* O-antigen ligase WaaL (Fig.1A & Supplementary Fig.1A) [24, 25]. The same band patterns were also visualised by commercially available Group A Carbohydrate specific antibodies, confirming detection of the correct carbohydrate epitope, in agreement with our previous work on the gene cluster [9]. We isolated OMVs from *E. coli* liquid cultures, which were either decorated with RhaPS (RhaPS-OMVs) (Fig. 1B) or lacked RhaPS (empty OMVs). When probed with GAC-specific antibodies, only the RhaPS-OMVs were detected, confirming the successful decoration of the OMVs with RhaPS (Fig. 1B). Before immunisation, the purified OMV’s were quantified and tested for LPS content using the LAL kit, revealing the endotoxin levels at 0.7 EU/ml (Supplementary Fig.1B). The OMV average particle sizes were in agreement with previously reported *E. coli* OMVs, with a diameter range of 50–250 nm (Supplementary Fig.1C) [26].

**Fig. 1:**
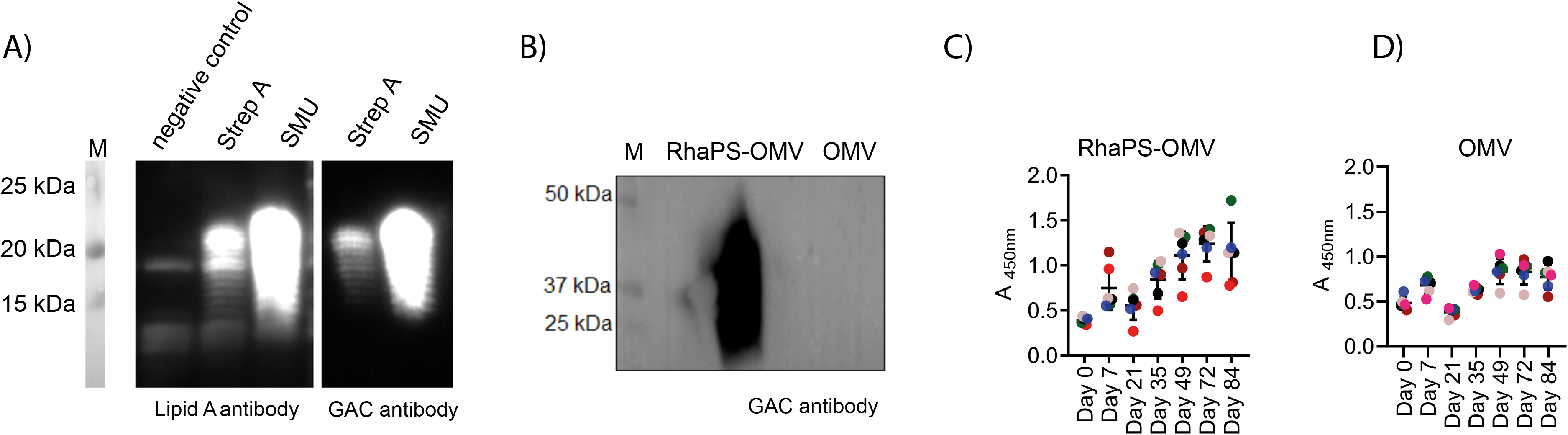
Purification and mice immunisation of recombinantly produced RhaPS-OMV: A) Western blot of RhaPS produced in *E. coli* cells from the *Streptococcus pyogenes* and *Streptococcus* mutans (SMU) gene clusters and probed with Lipid A antibody and GAC specific antibody. Negative control cells not producing the RhaPS cluster in lane (-). B) Representation of immunoblot analysis of recombinantly produced *E. coli* producing RhaPS-OMV and *E. coli* OMV were stained for 1:1000 Rabbit anti-GAC antibody followed by goat anti-rabbit IgG HRP. C&D) 96-well plates coated with *E. coli* cells expressing the RhaPS from Strep A were analysed using sandwich ELISA for RhaPS-OMV specific IgG antibodies in anti-RhaPS-OMV sera (C) or OMV vaccinated mice sera (D) collected at each stage of immunisation. Bound antibodies were detected using pre-titrated, biotin-conjugated antibody. Each coloured dot represents an immunised animal that has been tracked from day 0 to day 84. Data shown are mean±S.E.M. from technical replicates.

OMVs have been shown to induce IgG antibodies in animals [27] as well as in humans [28]. We immunised mice with either RhaPS-OMVs or control OMVs and measured the levels of RhaPS-OMV-specific IgG using an ELISA assay, targeting *E. coli* cells that either express or lack RhaPS on their outer membrane (Fig.1C, D). The RhaPS-OMV-specific IgG levels were significantly higher in RhaPS-OMV immunised animals compared to control animals (immunised with ‘empty’ OMVs), indicating that RhaPS-OMVs are more immunogenic than OMVs lacking the RhaPS carbohydrate (Fig. 1C, D). The presence of baseline IgG levels in the anti-OMV sera when probed in an ELISA assay with E. coli cells expressing RhaPS is not unexpected, as the surface of these cells contains elements common to both native OMVs and OMV-RhaPS.

### RhaPS-OMV stimulated antibodies selectively bind to the RhaPS epitope

Final bleed sera from each mice group were pooled and analysed by flow cytometry for antibodies that recognise the RhaPS antigen expressed on *E. coli* cells and compared with control cells lacking RhaPS. Histograms for antibody deposition on RhaPS positive *E. coli* cells or RhaPS negative *E. coli* cells labelled with the different antisera from RhaPS-OMV and OMV immunised mice reveals two distinct plots, suggesting that RhaPS specific antibodies are triggered that bind to the *E. coli* cells that produce RhaPS (Fig.2A). The extraction of geometric mean fluorescence intensity (gMFI) from the flow cytometry data clearly revealed increased IgG antibody deposition onto *E. coli* RhaPS-decorated cells, compared to control anti-OMV IgG (Fig.2B), indicating that selective paratopes are expressed in the sera of the RhaPS-OMV vaccinated animals. Importantly, the gMFI for RhaPS-OMVs were significantly higher than of the negative control OMVs on exposure to RhaPS positive *E. coli* cells, suggesting strong immunogenicity of the RhaPS (Fig.2B). These findings align with our observation that OMVs alone stimulated the production of antibodies that recognise *E. coli* cells without recombinantly produced RhaPS, and to a lesser extent, *E. coli* decorated with RhaPS, which shields the outer membrane (Fig. 2B). Interestingly, the levels of cross-reactivity were similar for both species (∼50 gMFI units), supporting our interpretation that RhaPS-OMVs stimulate production mainly of antibodies against the RhaPS but also, to a lower extent, against other components present in the OMVs.

**Fig. 2:**
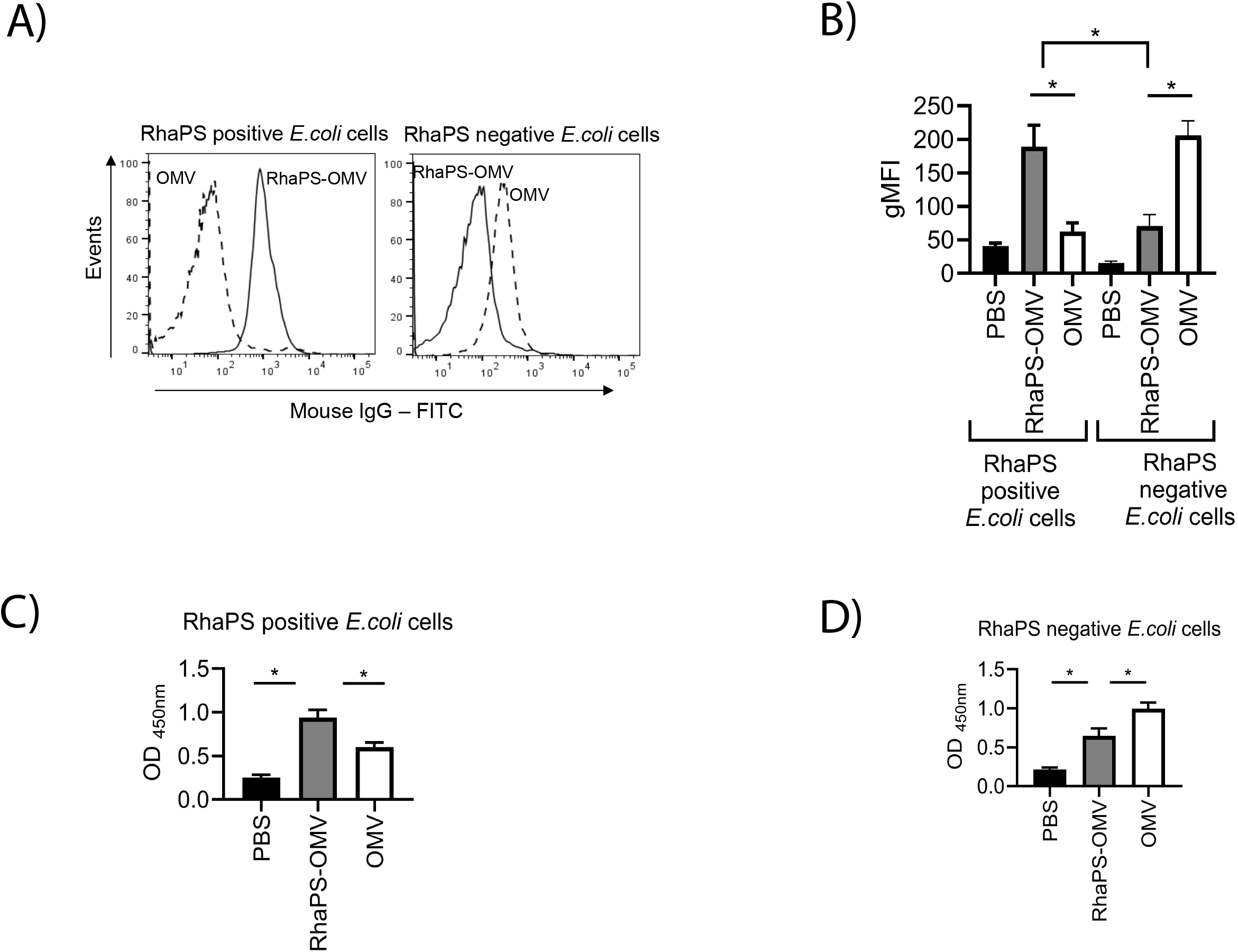
RhaPS-OMV induced IgG antibodies selectively binds to RhaPS positive *E. coli* cells: A) Representative histograms for antibody deposition on RhaPS positive *E. coli* cells or RhaPS negative *E. coli* cells stained either using day 84 final pooled antiserum (1:100) from RhaPS-OMV immunised sera (open black line) or anti-serum from OMV alone immunised sera (1:100) (dashed line). The stained cells were probed with anti-mouse IgG FITC and analysed in flow cytometry. (B) IgG deposition of RhaPS-OMV, OMV alone, or PBS (sham) immunised animal sera from day 84 analysed using geometric mean fluorescence intensity (gFMI) by treating either with the RhaPS positive *E. coli* cells or RhaPS negative *E. coli* cells. C,D) Plates were coated with C) RhaPS positive *E. coli* cells or D) RhaPS negative *E. coli* cells and probed with a 1:1,000 dilution of day 84 murine sera recovered from RhaPS-OMV, OMV, or PBS-sham immunised mice. Bound antibodies were detected using a 1:1,000 dilution of HRP-conjugated goat anti-mouse IgG. Flow data shown are collected from at least 10,000 events. Statistical analyses were conducted using ANOVA followed by Bonferroni post hoc test *p<0.05. Data shown are mean±S.E.M. of three independent experiments.

We developed an ELISA assay using *E. coli* cells and tested the post-immunisation sera from the mice in this secondary assay. Serum from the RhaPS-OMV-vaccinated animals exhibited significantly higher binding affinity for RhaPS-bound *E. coli* compared to serum from animals immunised with OMVs or PBS (Fig. 2C). These data confirmed that the RhaPS antigen on the *E. coli* cells were recognised robustly by antibodies stimulated in the RhaPS-OMV-immunised animals. In comparison, OMV-immunised animal sera showed significantly less signal (Fig.2C), and the anti-OMV titres in these animals were significantly higher in negative control cells alone compared to anti-RhaPS-OMV titre groups (Fig.2D). Immunoblot analysis of RhaPS-OMV immunised serum showed a strong signal in lysates from *E. coli* cells that produce RhaPS confirming the presence of RhaPS specific antibodies (Supplementary Fig.1D).

### Validation of IgG subtypes in RhaPS-OMV immunised mice sera

IgG subtypes have the ability to drive increased affinity for polysaccharide antigens in animals and humans [29-31]. Therefore, we examined the effect of RhaPS-OMV on the IgG subtypes in mice (IgG1, IgG2a, IgG2b and IgG3) using the Luminex analysis (Fig.3). The IgG subtypes were analysed in pooled sera of the PBS and RhaPS-OMV immunised animals on day 49 (after the administration of two boosters) and on day 70 (after three boosters) (Fig.3). Our results show that all tested IgG subtypes were significantly increased in RhaPS-OMV immunised animals, with the largest increases observed in IgG2a (Fig. 3B), IgG2b (Fig. 3C), and IgG3 (Fig. 3D), and a more modest increase in IgG1 levels (Fig. 3A). Significantly increased titres of anti-RhaPS-OMV IgG subtypes, compared to PBS immunoglobulin levels, were sustained until the end of the immunisation schedule (day 70). Our objective was to ultimately target Strep A bacteria and not *E. coli* cells, therefore, we did not include a group immunised with *E. coli* OMVs alone, which gave limited background antibodies in our pilot study against *E. coli* cells and OMVs (Fig. 1D). However, we acknowledge that including OMV sera would have provided additional insights into OMV-driven IgG subtype responses.

**Fig. 3:**
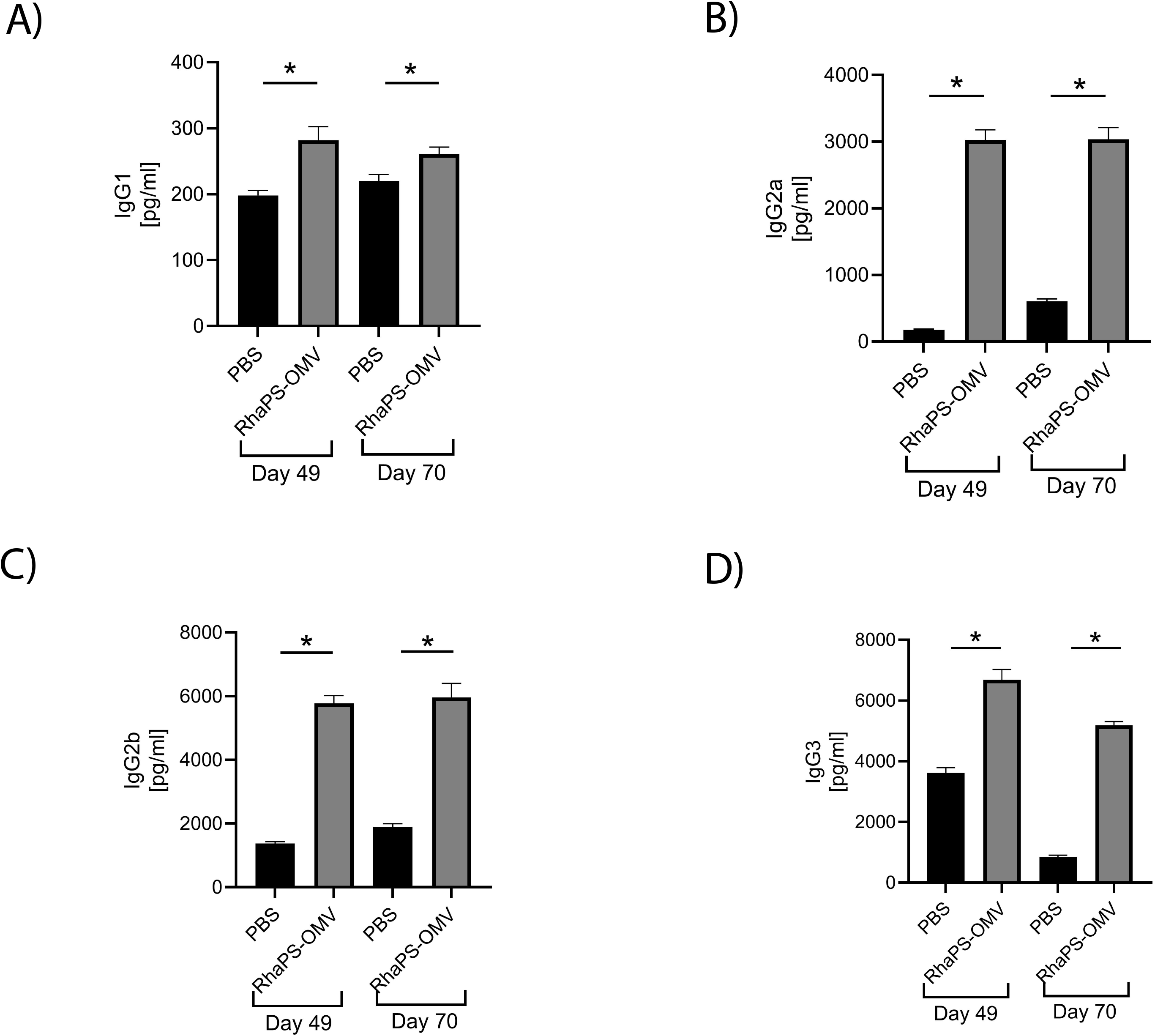
Quantification of Immunoglobulin subtypes against RhaPS-OMV in mice sera using Luminex analysis: A) IgG1, B) IgG2a, C) IgG2b, and D) IgG3 immunoglobulins were compared between day 49 (after two boosters) and day 70 (after three boosters) on the pooled RhaPS-OMV immunised mice sera or with PBS sera and analysed using Luminex. *p< 0.05 unpaired two-tailed T test (*vs*. PBS). Data shown are mean±S.E.M. from technical replicates.

### RhaPS-OMV antibodies recognise diverse Strep A serotypes

The efficacy of the RhaPS-OMV antibodies to opsonise to the universally conserved carbohydrate in a variety of Strep A isolates was investigated using flow cytometry and ELISA analyses. Strep A bacteria were exposed to pooled sera from the different groups and the gMFI obtained from the flow cytometry data were analysed. Significantly increased antibody deposition for RhaPS-OMV antibodies were detected on M1_5448_, M6, M11 and M89 bacteria when compared to OMV control immunised sera (Fig.4A). Furthermore, increased binding was observed for M2, M3, M75 and M1_UK_ (Fig.4A). However, M12 and M28 Strep A serotypes were not recognised by the RhaPS-OMV IgG. Whilst OMVs are considered naturally immunogenic and often do not require an additional adjuvant to enhance the immune response. However, considering that the immune response in mice was not as significant as expected, when compared to chemical conjugation approaches [11, 12]. We therefore conducted a second immunisation study where the OMV immunogens were supplemented with aluminium adjuvant. Post-immunisation sera were pooled and analysed for their ability to target Strep A bacteria. On average, opsonisation was two-fold stronger for all tested serotypes from the RhaPS-OMVs immunisation study, with only two out of ten tested Strep A strains not being significantly increased (Fig.4B). The degree of IgG binding to the Strep A isolates varied highly across all tested serotypes, in agreement with previous reports using chemical conjugated RhaPS-based vaccine candidates [11, 12]. In conclusion, RhaPS-OMV supplemented with aluminium provided increased opsonisation of Strep A bacteria (Fig.4A,B). A secondary ELISA assay confirmed that aluminium-adjuvanted RhaPS-OMV sera recognised all Strep A serotypes significantly better than the adjuvant control sera, including the dominant M1T1 clade (M1_UK_), a new *emm*1 *S. pyogenes* lineage (Fig.4C) [32].

**Fig. 4:**
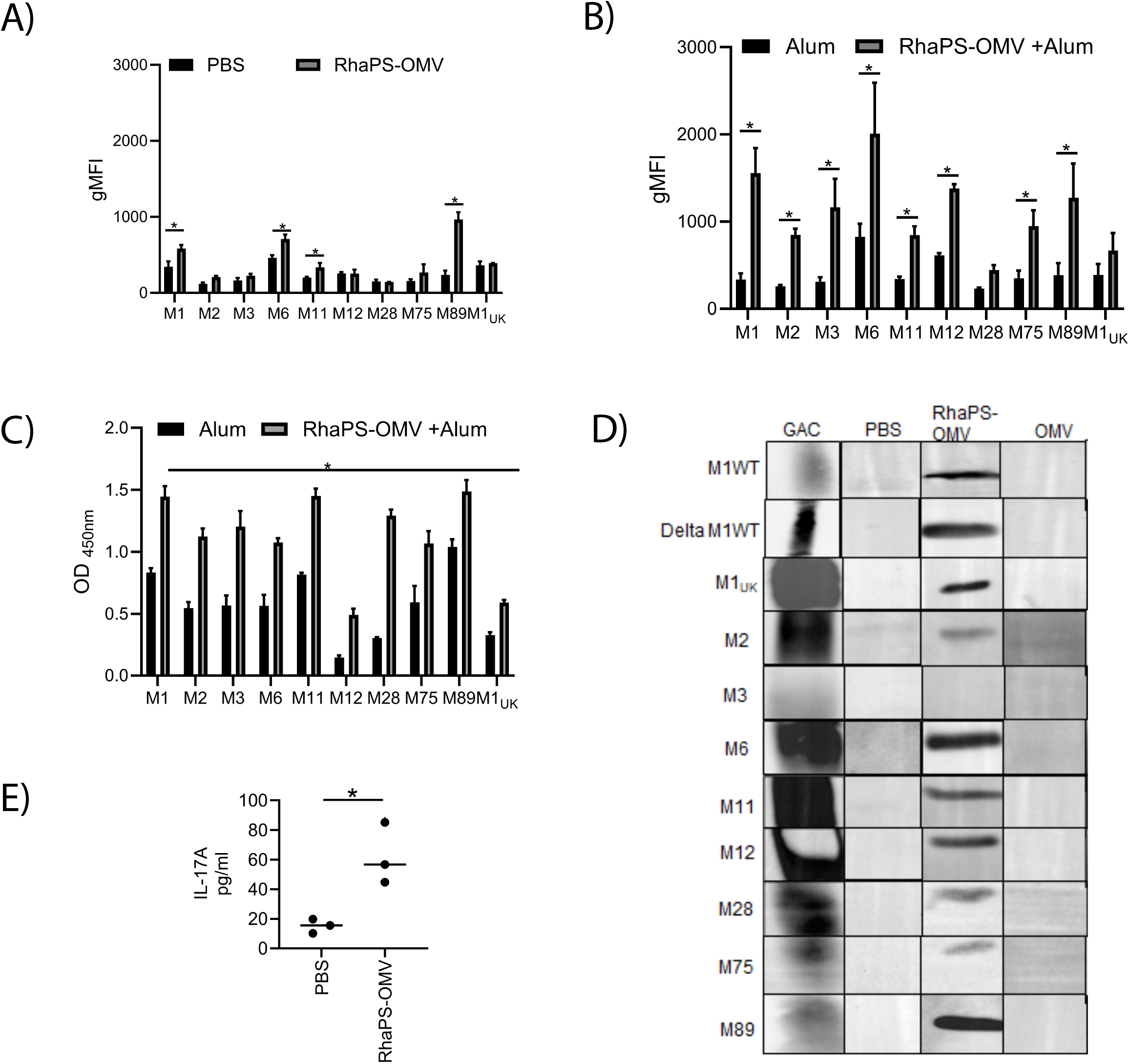
RhaPS-OMV vaccinated antibody deposition on Strep A isolates: A) Graph showing geometric mean fluorescence intensity (gMFI) extracted from flow cytometry analysis of clinical Strep A isolates stained with either PBS or RhaPS-OMV pooled mice sera from day 70 (1:100) followed by anti-mouse IgG AlexaFluor 488 channel B) Graph showing gMFI extracted from flow cytometry analysis of clinical Strep A isolates stained with either alum or RhaPS-OMV with alum pooled mice sera from day 8 (1:100) followed by anti-mouse IgG AlexaFluor 488 channel C) Whole cell ELISA analysis conducted to measure the antiserum (day 8) from RhaPS-OMV with alum (1:1,000) or antiserum from alum (1:1,000) binding to clinical Strep A strains D) Immunoblot analysis of clinical Strep A isolates stained using pooled mice sera from day 8 from the animals immunised with either PBS, RhaPS-OMV or OMV were used to probe the Strep A lysates at 1:1000 dilution. The Strep A lysates were further probed with anti-mouse IgG HRP (1:1,000). Anti-Group A Carbohydrate antibody (GAC) was used as a positive control (1:1,000). E) Graph showing the cytokine mediator IL-17a measured in the supernatants of individual mice splenocytes (PBS or RhaPS-OMV) restimulated with recombinant OMV RhaPS antigen (10 μg/ml). Results displayed as mean ± SEM from technical replicates. Statistical analyses were conducted using ANOVA followed by Bonferroni post hoc-test *p<0.05 (PBS *vs*. RhaPS-OMV or Alum vs RhaPS-OMV+Alum). Data shown are mean±S.E.M. of three independent experiments.

Next, we investigated if the mice antibodies were able to bind to bind to the Strep A serotypes in a Western blot assay, where protein samples are denatured. This analysis should provide more specific information whether or not the GAC carbohydrate was detected. Whole cells were inactivated, and the cell lysate separated *via* an SDS-PAGE. The lysates were then probed with different antibodies, including commercially available Group A Carbohydrate antibodies that detect the surface carbohydrate but also binds to other cell wall components/proteins. We compared this generic Strep A antibody with our RhaPS-OMV and negative controls (OMV alone or PBS-immunised) mice sera. Strep A isolates show an intense band at ∼25 kDa on Western blots probed with sera from RhaPS-OMV mice. The band was absent in the control OMV sera, confirming the specificity of the RhaPS antibodies to the native Strep A GAC carbohydrate. The positive control (commercial Strep A antibodies) displayed a stronger signal representing the presence of complete GAC components (including GlcNAc/glycerol-phosphate sidechain) and most likely other protein components (Fig.4D). As a control to investigate the specificity of the commercial Strep A antibodies, we included the strain Δ*gacI*, which produces only the RhaPS backbone [11]. The band pattern is equally broad, suggesting that other components in the lysate are recognised by the polyclonal antibodies. This is not surprising, considering that these antibodies were raised against heat-inactivated Strep A bacteria and not against the purified Group A Carbohydrate.

### RhaPS-OMV sera stimulate IL-17a production in murine splenocytes

We investigated the effect of the RhaPS-OMV immunogen on IL-17a production in splenocytes following immunisation. IL-17a, a key cytokine mediator, contributes to activation of T cells by mediating protective innate immunity against Strep A [33]. Increased IL-17a levels were detected in RhaPS-OMV vaccinated splenocytes restimulated with RhaPS-OMV antigen, compared with splenocytes from the PBS group (Fig.4E). This is the first study that suggests that RhaPS-OMV induces a humoral mediated immune response, and also a cytokine-mediated cellular immune response [33, 34], potentially via carbohydrate-specific T helper cells [35].

### RhaPS-OMV immunised rabbit serum bind to Strep A bacteria from *S. pyogenes* and *S. dysgalactiae* subsp. *equisimilis*

We next studied whether RhaPS-OMV immunisation in rabbits triggers antibody responses similar to those observed in mice. Post-immunisation sera were first tested in an immunoblot using *E. coli* lysates for the presence of anti-RhaPS antibodies (Fig.5A). Only sera from RhaPS-OMV immunised rabbits were able to detect the recombinantly produced carbohydrate, consistent with the results obtained using commercially available GAC antibodies. Post-immune raw sera and IgG antibodies also recognised several potential OMV proteins, which are present in both the RhaPS-OMVs and the OMV-negative control immunogens, and thus were also recognised by sera from rabbits immunised with OMV controls. This is not surprising, given that the OMVs contain common E. coli proteins, which are capable of stimulating an immune response. Next, we utilised an ELISA to investigate the pre- and post-immunisation sera for reactivity against a RhaPS-glycoconjugate protein, which consists of a carrier protein conjugated with the Strep A polysaccharides (NanA-RhaPS) [36]. The ELISA confirmed that all three rabbits produced IgG antibodies against the RhaPS carbohydrate, independent of the *E. coli* OMV specific proteins (Fig.5B). The successful immunisation of RhaPS specific antibodies in rabbits led us to test the antibodies’ ability to opsonise various Strep A serotypes, following the above reported approach with mice sera (Fig.5C). The histograms revealed that rabbits vaccinated with RhaPS-OMV exhibited a stronger reaction to all tested Strep A serotypes compared to their pre-immune sera. This is indicated by higher gMFI values and a more intense blue shade for FITC-positive cells, demonstrating a stronger response relative to the pre-vaccination sera (Fig. 5C and Supplementary Fig. 2).

**Fig. 5:**
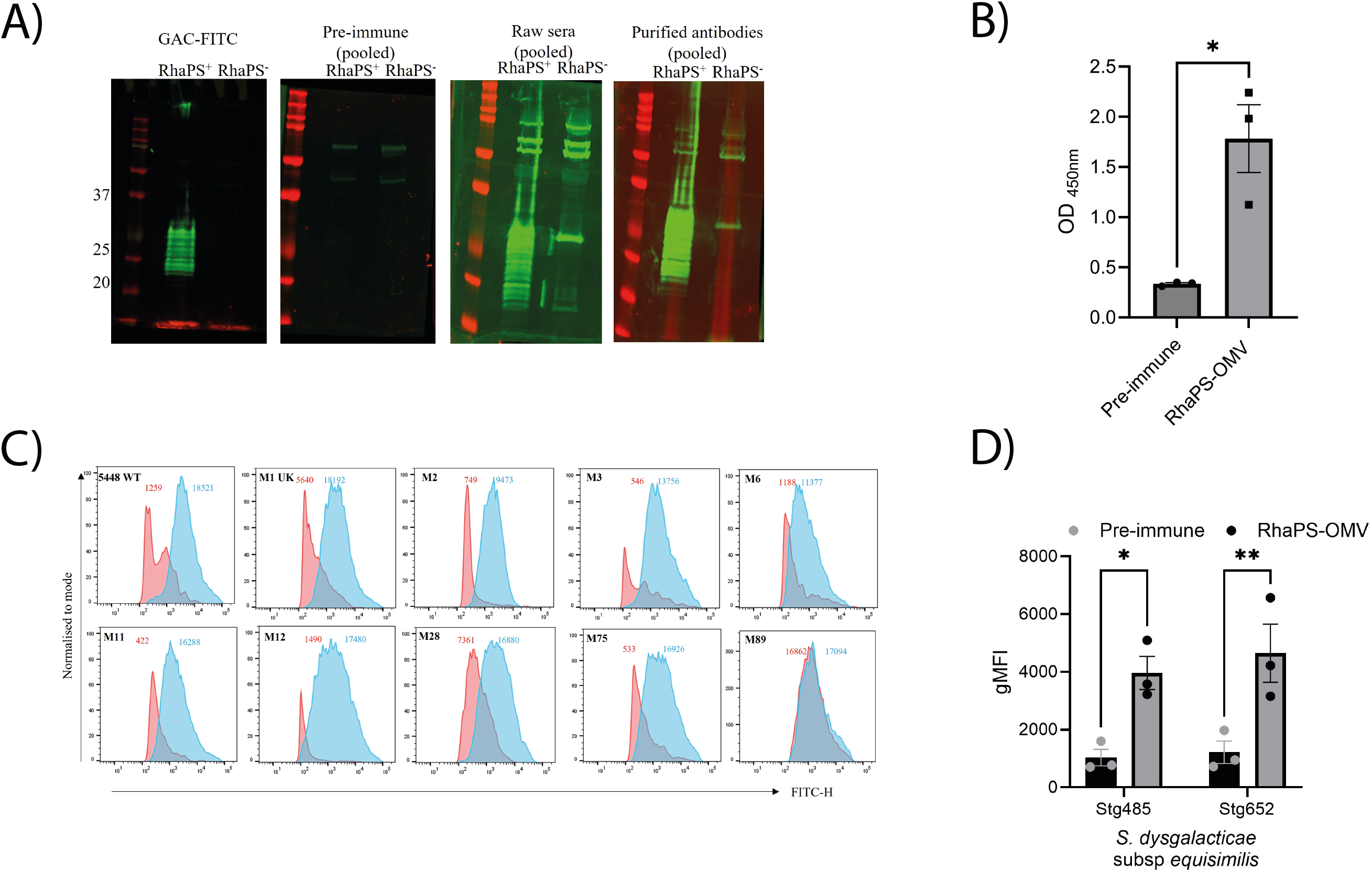
Rabbit immunised RhaPS-OMV IgG promotes binding to Strep A M strains and *Streptococcus dysgalactiae* subsp. *equisimilis* isolates. A) Immunoblot analysis of *E. coli* cells expressing RhaPS were measured by using the pooled antiserum from the immunised OMV-RhaPS (raw sera) or affinity purified OMV-RhaPS or the pre-immune sera (1:2,000). Molecular mass markers are given in kilodaltons. B) Anti-OMV-RhaPS IgG antibodies were measured by ELISA using individual rabbit pre-immune and OMV-RhaPS immunised sera (1:1,000) from day 63. The ELISA plate were coated with a glycoconjugate protein (NanA-RhaPS) at 20 μg/ml. Data displayed are mean ± SEM from three individual rabbit sera. Unpaired T test analyses (*p<0.05) were performed using GraphPad Prism. C) Representative histograms for antibody deposition on Strep A M strains using individual rabbit antiserum (1:1000) for all ten strains. Pre-immune is represented as red shading and post-immune OMV-RhaPS sera is represented in blue shading. The number on the top shows the geometric mean fluorescence intensity (gMFI) of the pre-immune sera (red) and OMV-RhaPS immunised serum (blue) for the displayed histograms. D) Antibody deposition measured using a flow cytometry assay on *Streptococcus dysgalacticae* subsp. *equisimilis* containing GAC and lacking Group G Carbohydrate (GGC) (Stg485 and Stg652) stained in 1:1,000 rabbit antiserum from pre-rabbit sera and RhaPS-OMV immunised sera.

Having established that RhaPS-OMV antibodies raised in rabbits are efficient in binding Strep A isolates from *S. pyogenes*, we investigated the effect of these sera to also target the newly emerged *S. dysgalactiae* subsp. *equisimilis* (SDSE) isolates, which have replaced their Group G Carbohydrate gene cluster with a functional Group A Carbohydrate cluster (SDSE_*gac*) [21]. SDSE_gac isolates Stg485 and Stg652 were investigated in our flow cytometry assay. The calculated gFMI verified that antibodies in the post-immune RhaPS-OMV sera recognised and bound to these SDSE_*gac* isolates (Fig.5D) confirming that the GAC on SDSE isolates can also be targeted with RhaPS specific antibodies. These results highlight the additional advantage of a RhaPS-based vaccine, which could also be effective against these and other newly emerging streptococcal pathogens.

Microscopic examination of a representative Strep A serotype (M89) using RhaPS-OMV rabbit serum paired with fluorescently labelled anti-rabbit secondary antibodies displayed uniform cocci (long chains) revealing the classical structure of Strep A (Supplementary Fig.3A). Contrary the pre-immune serum showed only background fluorescent signal, further supporting our findings that recombinantly produced RhaPS in OMV triggers antibodies targeting the surface of Strep A bacteria and providing a valuable universal vaccine candidate packaged into outer membrane vesicles.

## Discussion

Several recent studies have addressed the production of universal Strep A vaccine candidates targeting the Strep A RhaPS backbone or the GlcNAc-sidechain decorated version of the Group A Carbohydrate (reviewed in [17, 36, 37]). These approaches focused on chemical/enzymatic extraction of natively produced RhaPS from Strep A cells, chemical synthesis of short GAC repeat units, followed by chemical conjugation to a carrier protein or OMV [12, 38, 39]. We explored an alternative production route and, for the first time, successfully produced recombinant RhaPS embedded in *E. coli* OMVs, evaluating them as vaccine candidates against Strep A. OMV vaccines are considered a promising approach in vaccine development, utilising genetically modified bacteria to produce outer membrane vesicles decorated with pathogen-specific carbohydrates.

The data demonstrate that RhaPS-OMVs were effectively presented to the animals’ immune systems and, when supplemented with the commonly used and safe alum adjuvant, triggered a strong immune response specific for the RhaPS carbohydrate. This effect agrees with the findings of a study investigating *E. coli* OMV decorated with β-(1→6)–linked poly-N-acetyl-D-glucosamine (PNAG) [40]. PNAG is a conserved polysaccharide antigenic component found in bacteria, fungi, and protozoan cells. Immunisation studies on PNAG-OMV provoked polysaccharide specific IgG antibodies.

The prevalence of IgG subtypes has been widely studied within the Strep A infection models. One of the most abundant subclasses of human IgG found against the Strep A M proteins is IgG3 [41]. Notably, antibody isotype class switching from IgM to mouse IgG3 constant regions was proven to be driven by the frequent exposure to polysaccharide antigens [42, 43]. A strong IgG1 response was observed in humans exposed to polysaccharides from pneumococcal pathogens [29] and increased IgG2a and IgG2b levels were detected in mice exposed to M protein-based vaccines from *Streptococcus pyogenes* [44].

Considering recent evolutionary events that resulted in antigen exchange from Group G/C streptococci into Strep A bacteria, it is important to develop a vaccine that also has the ability to target these emerging strains. The successful recognition of SDSE_*gac* isolates by RhaPS-OMV antibodies therefore broadens the potential protection of a RhaPS-based vaccine to target newly emerging SDSE isolates.

A major finding of this study is the increased expression of IL-17a stimulated by the RhaPS-OMV antigen in mouse splenocytes. Carbohydrates are known to trigger a T cell independent immune response and hence conjugating them with a carrier protein often elicits T cell dependent immune response. It has been demonstrated that the recognition of polysaccharides by CD4+ T cells triggers their activation and differentiation, which can be assessed by measuring the IL-17A cytokine. [45]. IL-17-mediated protective immunity via neutrophil recruitment has been shown against Strep A [33], *Klebsiella pneumoniae* [46] and *Streptococcus pneumoniae* [47]. However, we cannot rule out the possibility of a role for endogenous *E. coli* antigens present in the OMV that are able to induce IL-17a, which needs further exploration.

In recent years, several OMV vaccines have been licensed against bacterial pathogens, including *Neisseria meningitis* [48] with other OMV based vaccines being in clinical trials [49, 50]. The technology offers advantages in terms of vaccine development speed, production scalability, and cost-effectiveness, making it an attractive option for addressing global health challenges, especially in low- and middle-income countries. Our approach, paired for instance with hyper-vesiculating *E. coli* bacteria and strains that produce a reduced OMV proteome [51, 52] provides an appealing approach to develop an affordable glyco-conjugate vaccine targeting Strep A bacteria.

In summary, we have demonstrated that recombinantly produced RhaPS in E. coli OMVs induces robust IgG antibodies in mice and rabbits, specifically binding to all tested *S. pyogenes* and *S. dysgalactiae* subsp. *equisimilis* Strep A serotypes. The immune responses were stronger when RhaPS-OMVs were adjuvanted with alum. While several OMV vaccine candidates are reported to possess intrinsic self-adjuvant properties, the underlying mechanisms remain unknown [53]. At this stage, we can only speculate that variations in the effectiveness of self-adjuvanticity among different OMV-based vaccine candidates may depend on factors such as vesicle size, composition, and the specific carbohydrates coating the OMVs. Our RhaPS-OMV demonstrated greatly increased immunogenicity when administered with alum, an approved and safe adjuvant.

Importantly, we have shown for the first time that a Strep A vaccine candidate also has the ability to target the newly emerged GAC producing isolates from *S. dysgalactiae* subsp. *equisimilis*, which are also susceptible to RhaPS-OMV antibodies. This provides a valuable new opportunity for a Strep A specific vaccine that does not target strain specific proteins, but also targets other related pathogenic bacteria that produce the same Group A Carbohydrate structure. It remains to be evaluated if RhaPS-OMV also trigger IgG-mediated protection in *in vivo* Strep A infection models and how RhaPS-OMV efficacy compared against other Strep A vaccine candidates, including the recently reported chemical conjugation of recombinant produced RhaPS to a carrier protein [39].

## Methods

### Bacterial strains and growth conditions

A single *E. coli* colony was cultured in 10 mL LB medium supplemented with 150 µg/mL erythromycin, grown at 37°C, 200 rpm overnight and used to inoculate 1 L medium, grown for 18 h. Strep A isolates were grown at 37 °C, 5% CO_2_ in THY. *E. coli* and Strep A bacteria were used as overnight cultures for Western blot analysis or used at 1:100 to O.D_600_ 0.4 for flow cytometry and ELISA assays.

### Purification of RhaPS-OMVs

*E. coli* OMVs were isolated from CS2775 *E. coli* cells transformed with plasmids encoding the RhaPS backbone biosynthesis genes (pHD0136) and empty plasmid control (pHD0139), respectively [9]. Cells were harvested by centrifugation (4,500 RPM, 30 min, 4ºC). The supernatant was filtered (0.45μm) and centrifuged (Type 45Ti, 45,000 RPM, 4 hrs, 4ºC) to isolate the OMVs. The pellets were weighed, resuspended in PBS, and protein content determined using the Bradford method. The OMVs were analysed for their average size distribution in the Zetaview nanoparticle tracking analysis instrument, and LPS levels assessed using the Limulus Amoebocyte Lysate (LAL) assay (Thermo Fisher, UK).

### Immunisation

Mouse Model: Five-to six-week-old female C57BL/6J mice were acquired from Charles River Laboratories, UK, and were acclimatised for 10 days prior to immunisation. Mice (n=6) were immunised subcutaneously with RhaPS-OMV (10 µg diluted in 100 μl of endotoxin free PBS/mouse/injection) or *E. coli* OMV (n=6) on day 0, immediately following tail bleed for pre-immune sera. Identical booster injections were administered on day 21 and day 49. Animals were euthanised by CO_2_ asphyxiation and bled by cardiac puncture on either day 70 or day 84. For adjuvant studies, Imject Alum (Sigma) was used with 10 µg of RhaPS-OMV (1:1 ratio); adjuvant with PBS only (1:1 ratio) served as a negative control. PBS/Alum vaccinated mice (n=3) were used for baseline measurements.

Rabbit Model: For New Zealand White rabbit immunisation, the animal (n=3) was bled for pre-immune serum prior to immunising with the purified OMV RhaPS (150 µg diluted in 150 μl of endotoxin-free PBS/injection) on day 0. Booster injections were administered on days 14, 28, 42 and 56, at 100 µg in 100 μl of endotoxin-free PBS/injection. The animal was culled by CO_2_ asphyxiation and bled by cardiac puncture. Final bleed serum was taken on day 63 and affinity-purified (0.53 mg/ml) by Davids Biotechnologie GmbH.

### Flow cytometry

Antibody binding to Strep A and *E. coli* cells was adapted from [54]. Bacterial cells (2 × 10^6^ CFU) were resuspended in nonspecific human IgG (Sigma, UK) for one hour on ice, then washed with PBS and diluted 1:100 with either immunised mice or rabbit sera (PBS or RhaPS-OMV) overnight at 4°C. Cells were washed twice with PBS and stained with 1:250 dilution of Alexa fluor 488-conjugated goat anti-mouse IgG (Thermo Fisher, UK) or goat anti-rabbit FITC (Invitrogen), incubated for 20 minutes at 4°C in the dark. Cells were washed twice (PBS) and fixed with 500 μl of 4% PFA and analysed by flow cytometry (BD Bioscience). A total of 10,000 events were acquired and data were analysed using FlowJo software version 10.6.2.

### ELISA

For antigen-antibody interaction, plates were coated with 50 μl/well of 20 μg/ml OMV-RhaPS/negative control OMVs or *S. pyogenes* cells adapted from [55, 56]. Briefly, bacterial cells at OD_600_ of 0.4, from the overnight culture, were resuspended (1:100) in PBS. Mouse or rabbit serum from immunised animals was added, 50 µl/well at 1:1,000. Bound antibodies were probed using 50 μl/well of 1:1,000 of anti-mouse IgG HRP (Sigma–Aldrich) and 75 μl/well of tetramethylbenzidine substrate (Sigma–Aldrich). The reaction was stopped using 75 μl/well of 1 M H_2_SO_4,_ and absorbance read at 450 nm.

### Luminex

Immunoglobulins in mouse sera were quantified using a Milliplex Multiplex assay (Merck Millipore, UK). Immunoglobulin beads (IgA, IgG1, IgG2a, IgG2b) from Mouse Antibody Isotyping 7-Plex ProcartaPlex were used as analytes and the samples were prepared according to manufacturer’s instructions (Thermo Fisher, UK).

### Immunoblot analyses

Antisera from RhaPS-OMV vaccinated animals were tested for binding to *E. coli* cells or Strep A by Western blot analysis using 12.5% acrylamide PAGE gels (Thermo Fisher, UK). *E. coli* cells and OMVs were used from overnight cultures. Overnight grown Strep A cells were washed with PBS and incubated with 6 μl of PlyC (0.7 mg/ml) for one hour (37°C, 300 RPM) and centrifuged for 14,000 RPM, 5 mins. The resulting pellets were resuspended in 2x SDS-PAGE loading dye and subjected to Western blot analysis (Invitrogen, UK). The PVDF membranes were blocked with 5% non-fat dried milk in Tris-Buffered Saline, 0.1% Tween® 20. Immunised mouse sera were used at 1:1000 dilution to probe the blots (overnight, 4°C, 20 RPM) followed by goat anti-mouse IgG HRP at 1:1000 dilution for two hours at 4°C, 20 RPM. Rabbit anti-GAC antibodies (Abcam, UK) with goat anti-rabbit IgG HRP (Abcam, UK) (1:2,500) were used as a positive control. Antibody binding was visualised using Clarity Western ECL Substrate (Biorad, UK) and viewed under the Gel Doc imaging system (GeneSys software, Syngene).

### Isolation and *ex vivo* re-stimulation of splenocytes

Isolation of mouse spleens and re-stimulation with antigens was conducted according to published procedures [57]. Briefly, spleens were rinsed and flushed, using 21G needles, with 5 ml of RPMI 1640 media (supplemented with 100 U/ml Pen-Strep (Sigma, UK), 200 mM L-glutamine (Gibco, UK) and 10% heat-inactivated foetal bovine serum (Sigma, UK)). The dispersed cells were centrifuged (300 x *g*, 10 min) and red blood cells were lysed in lysis buffer (Sigma, UK). Cells were washed and centrifuged again by the addition of 10 ml of PBS (300 x *g*, 10 min). Pellets were resuspended in complete RPMI 1640 media and adjusted to 7.5 × 10^6^ cells/ml; 1.5 × 10^6^ cells [200 µl] were added to each well of a 96 well plate followed by 50 µl of 1 µg/ml RhaPS-OMV and incubated for three days. Cells were pelleted (400 x *g*, 10 min) and supernatants subjected to ELISA analysis for IL-17a (Thermo Fisher, UK).

### Microscopic analysis

Strep A M89 cells were grown as mentioned above. Briefly, overnight culture of M89 strains were washed twice with PBS (10,000 RPM, 5 minutes). Washed cells were stained overnight at 4°C with pre-rabbit or RhaPS-OMV post immune sera at 1:100 dilution. Prior to adding secondary antibody (goat anti-rabbit FITC in 1:50) the cells were washed twice with PBS. The FITC-stained cells were mounted and viewed using a DeltaVision microscope; images were deconvoluted and analysed using softWoRx imaging system.

### Ethical approval

Mice studies were approved by the University of Dundee welfare and ethical use of animals committee and conducted in accordance with the UK Home Office approved project licence [PPL PEA2606D2]. The experiments involving rabbits were conducted in association with Davids Biotechnologie GmbH, Germany.

### Statistical Analysis

Data were analysed using GraphPad Prism Version 8 (La Jolla, CA, USA). One-way analysis of variance [ANOVA] with Bonferroni’s correction, two-tailed T-test followed by Dunn’s post-test were used to test the statistical significance. FlowJo analysis software (FlowJo, USA) was used to plot the geometric mean fluorescence intensity (gMFI) of flow cytometric data.

## Data availability

No datasets were generated or analysed during the current study.

## FIGURE LEGENDS

**Supplementary Fig.1:** A) Spot-Blot of the crude membrane (M), inner membrane (IM), and outer membrane (OM) of wildtype (WT) or ΔwaaL *E. coli* cells expressing the Strep A and SMU gene clusters probed with an anti-GAC antibody. B) Quantification of endotoxin levels in the purified RhaPS-OMV and OMV alone in first three doses used to immunise the animals. Number indicates the immunisations doses. C) OMV concentration and the particle size in diameter (nm) from the purified *E. coli* producing RhaPS or *E. coli* OMV on their own were estimated using ZetaView nanoparticle tracking analyser with 670 nm laser. Samples were diluted in 1:4 using PBS. PBS was used as a control and subtracted from all samples. Data shown are mean ± SD of technical replicates. D) Immunoblot analysis of whole *E. coli* RhaPS lysate (1) or *E. coli* empty plasmid control (2) were blotted using pooled antiserum from the RhaPS-OMV and OMV or PBS vaccinated groups. Bound antibodies were probed using a secondary antibody of 1:2500 dilution of anti-rabbit IgG HRP and viewed using image lab software. Molecular mass markers are given in kilodaltons (kDa).

**Supplementary Fig.2**: FITC positive cells were measured using flow cytometry assay on Strep A M strains in 1:1000 dilution of antiserum from pre-immune and OMV-RhaPS immunised sera. Paired T test analyses (*p<0.05, **p<0.01, ***p<0.001, ****p<0.0001) were performed using GraphPad Prism. Results displayed present mean ± SEM from individual rabbit sera (n=3).

**Supplementary Fig.3**: **Rabbit immunised RhaPS-OMV IgG promotes killing of the hypervirulent strain M89**: A) Representative images of immunofluorescent staining of hypervirulent strains M89 stained with either with pre-immune rabbit sera or RhaPS-OMV immunised rabbit sera (1:1,000) followed by goat anti-Rabbit IgG FITC (1:50) channel. Polarised (POL) and FITC channel were used to document the images using Deltavision microscopy. B) Percentage of survival of M89 hypervirulent Strep A strains were analysed using antiserum from the RhaPS-OMV immunised or pre-immune rabbit in the presence of 5% baby rabbit serum. *p < 0.05 unpaired two-tailed T test (PBS vs RhaPS-OMV or pre-immune sera vs post-immune sera). The data presented represent the mean ± S.E.M. from three independent experiments for B.

## Financial support

The H.C.D. laboratory was supported by the University of Dundee Wellcome Trust Funds 105606/Z/14/Z, Tenovus Scotland Large Research Grant [T17/17], Wellcome and Royal Society Grant 109357/Z/15/Z and Wellcome Innovator Award 221589/Z/20/Z. For the purpose of open access, the authors have applied a CC BY public copyright licence to any Author Accepted Manuscript version arising from this submission. H.A.S. is funded by Wellcome Innovator Award 221589/Z/20/Z and independent research commissioned and funded by the NIHR Policy Re-search Programme (NIBSC Regulatory Science Research Unit).

## Conflicts of Interest Statement

All authors declare no conflicts of interest.

## Acknowledgments

We thank Dr. Fatme Mawas for fruitful discussions.

## Author Contributions

S.A.C, S.T., H.A.S., A.Z. and B.H.M., performed the experiments, collected the data, and conducted the analysis. M.R. and H.C.D. analysed and interpreted data. S.A.C and H.C.D. wrote the manuscript. H.C.D supervised the study. All authors read and approved the final manuscript.

## Notes

### Competing Interest Statement

The authors have declared no competing interest.

