## Supplemental Figures 1-3 for "Rhamnose Polysaccharide-Decorated Outer Membrane Vesicles as a Vaccine Candidate Targeting Group A Streptococcus from *Streptococcus pyogenes* and *Streptococcus dysgalactiae* subsp. *equisimilis*"

### Supplementary Fig.1)

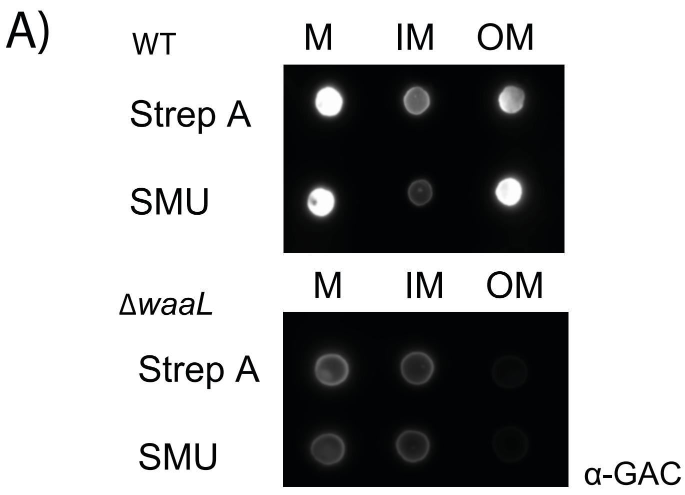


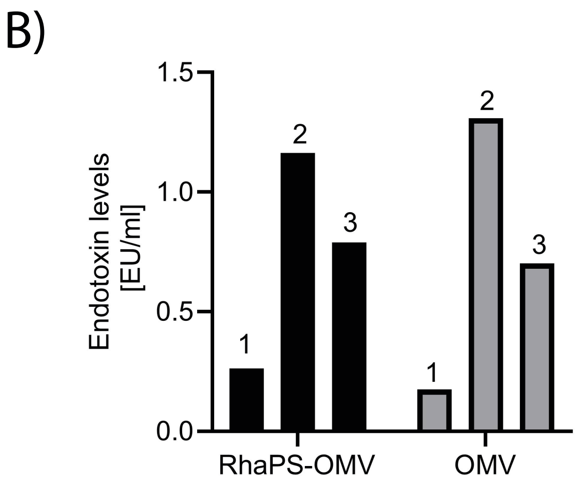


C)

| Sample | Count / mL | Average size |
|---|---|---|
| RhaPS-OMV | 4.0 e[9] | 131 nm |
| OMV | 3.0 e[7] | 153 nm |

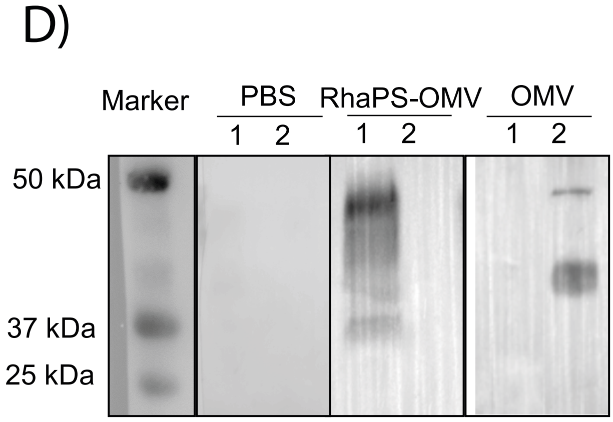


**Supplementary Fig.1:** A) Spot-Blot of the crude membrane (M), inner membrane (IM), and outer membrane (OM) of wildtype (WT) or ΔwaaL *E. coli* cells expressing the Strep A and SMU gene clusters probed with an anti-GAC antibody. B) Quantification of endotoxin levels in the purified RhaPS-OMV and OMV alone in first three doses used to immunise the animals. Number indicates the immunisations doses. C) OMV concentration and the particle size in diameter (nm) from the purified *E. coli* producing RhaPS or *E. coli* OMV on their own were estimated using ZetaView nanoparticle tracking analyser with 670 nm laser. Samples were diluted in 1:4 using PBS. PBS was used as a control and subtracted from all samples. Data shown are mean ± SD of technical replicates. D) Immunoblot analysis of whole *E. coli* RhaPS lysate (1) or *E. coli* empty plasmid control (2) were blotted using pooled antiserum from the RhaPS-OMV and OMV or PBS vaccinated groups. Bound antibodies were probed using a secondary antibody of 1:2500 dilution of anti-rabbit IgG HRP and viewed using image lab software. Molecular mass markers are given in kilodaltons.

### Supplementary Fig.2)

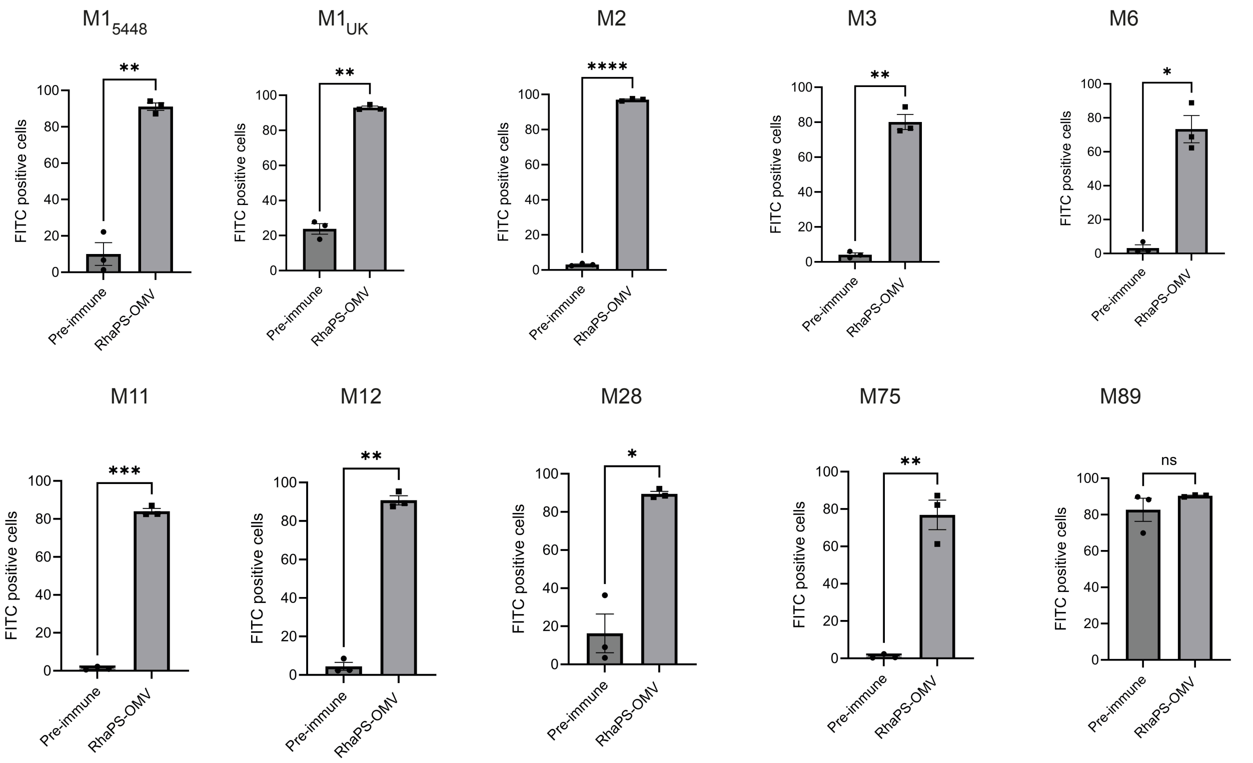


**Supplementary Fig.2**: FITC positive cells were measured using flow cytometry assay on Strep A M strains in 1:1000 dilution of antiserum from pre-immune and OMV-RhaPS immunised sera. Paired T test analyses (*p<0.05, **p<0.01, ***p<0.001, ****p<0.0001) were performed using GraphPad Prism. Results displayed as mean ± SEM from individual rabbit sera (n=3).

### Supplementary Fig.3)

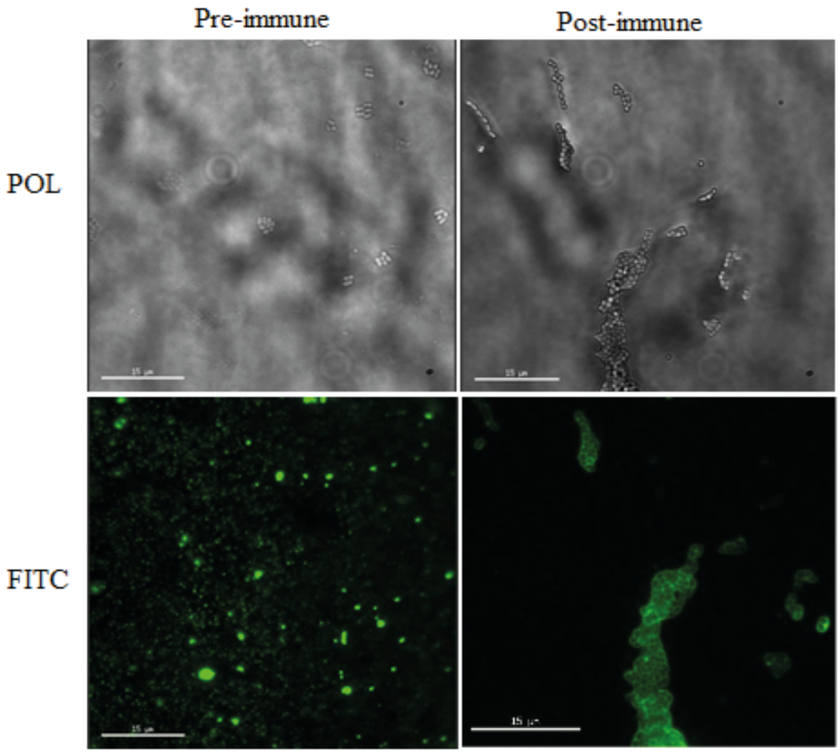


**Supplementary Fig.3**: **Immunoflurosecent staining of the hypervirulent strain M89 using antiserum from mice vaccinated with RhaPS-OMV IgG**: A) Representative images of immunofluorescent staining of hypervirulent strains M89 stained with either with pre-immune rabbit sera or RhaPS-OMV immunised rabbit sera (1:1,000) followed by goat anti-Rabbit IgG FITC (1:50) channel. Polarised (POL) and FITC channel were used to document the images using deltavision microscopy.
